# A transcallosal fiber system between homotopic inferior frontal regions supports complex linguistic processing

**DOI:** 10.1101/079244

**Authors:** Philipp Kellmeyer, Magnus-Sebastian Vry, Tonio Ball

## Abstract

Inferior frontal regions in the left and right hemisphere support different aspects of language processing. In the canonical model, left inferior frontal regions are mostly involved in processing based on phonological, syntactic and semantic features of language, whereas the right inferior frontal regions process paralinguistic aspects like affective prosody.

Using diffusion tensor imaging (DTI) based probabilistic fiber tracking in 20 healthy volunteers, we identify a callosal fiber system connecting left and right inferior frontal regions that are involved in linguistic processing of varying complexity. Anatomically, we show that the interhemispheric fibers are highly aligned and distributed along a rostral to caudal gradient in the body and genu of the corpus callosum to connect homotopic inferior frontal regions.

In light of converging data, taking previous DTI-based tracking studies and clinical case studies into account, our findings suggest that the right inferior frontal cortex not only processes paralinguistic aspects of language (such as affective prosody), as purported by the canonical model, but also supports the computation of linguistic aspects of varying complexity in the human brain. Our model may explain patterns of right hemispheric contribution to stroke recovery as well as disorders of prosodic processing. Beyond language-related brain function, we discuss how interspecies differences in interhemispheric connectivity and fiber density, including the system we described here, may also explain differences in transcallosal information transfer and cognitive abilities across different mammalian species.

## Introduction

Cumulative and converging evidence from clinical neurology, neuroanatomy, neuropsychology, and neuroimaging shows that the left and the right hemispheres support common as well as differing aspects of language processing. In this canonical model, the left perisylvian cortex mainly supports linguistic aspects of language processing like the analysis of phonological, syntactic or semantic features (Vigneau *et al.*, 2006; Hillis, 2007; Shalom & Poeppel, 2008; Price, 2012) and the right perisylvian cortex computes predominantly paralinguistic features of language like rhythm or prosody (Blonder *et al.*, 1991; Ross *et al.*, 1997; Frühholz *et al.*, 2015; Sammler *et al.*, 2015).

Macroanatomically, the left and right inferior frontal cortex (IFC) both consist of three parts: a superior dorsal part (pars opercularis and parts of ventral premotor cortex, IFC_po/PMv_), a middle part (anterior pars opercularis and posterior pars triangularis, LIFC_po/pt_) and an anterior-inferior part (pars triangularis, IFC_pt_). For cytoarchitecture, anatomists have found differences in volume and cell-packing density between left and right inferior frontal cortex for BA 44 and to a lesser extent for BA 45 (Amunts *et al.*, 1999). This suggests that BA 44 may be more left lateralized than BA 45 cytoarchitectonically.

For the role of left inferior frontal cortex in speech processing, aggregated evidence from twenty-five years of neuroimaging studies suggests a distinct function-anatomical organization in which the anatomical tripartition reflects a functional processing gradient (Bookheimer, 2002; Vigneau *et al.*, 2006; Price, 2010; Poeppel *et al.*, 2012; Kellmeyer *et al.*, 2013). In this model, the LIFC_po/PMv_ part is preferentially involved in phonological, the LIFC_po/pt_ in syntactic and the LIFG_pt_ part in semantic processing (Dapretto & Bookheimer, 1999; Shalom & Poeppel, 2008). Other neurolinguists have argued for a more supramodal view of left inferior frontal cortex in integrating syntactic and semantic features not exclusive to speech processing (Bornkessel-Schlesewsky & Schlesewsky, 2013).

The role of the homotopic right inferior frontal cortex in speech processing, however, is much less clear. In the canonical model described above, the right IFC mostly supports the analysis of paralinguistic, (specifically affective) features of prosody (Heilman *et al.*, 1984; Ross, 1993). Prosody, was originally proposed by the Norwegian neurologist Monrad-Krohn – the founder of the modern patholinguistic study of disorders of prosody – to be the “third element” of speech, the other elements being “grammar” (syntax) and “vocabulary” (semantics) (Monrad-Krohn, 1947, 1957). In this model, prosodic information is also conveyed by so-called *intrinsic* features like stress, rhythm and pitch and these features are processed by right *and* left inferior frontal cortex (Vigneau *et al.*, 2006, 2011; Frühholz *et al.*, 2015). Thus, the dynamic interaction between left and right inferior frontal cortex seems to be a necessary condition for successful language use in real-time. Psycholinguistic research with patients after corpus callostomy (usually for intractable epilepsy) in the 1970s has described processing deficits at the phonetic and semantic level, supporting this model of transcallosal integration (Levy & Trevarthen, 1977). These early findings, however, were not systematically explored and henceforth largely forgotten. Furthermore, it is difficult to ascertain to which degree the large-scale organization of the language system in patients with severe epilepsy differs from the healthy brain.

The left and right homotopic inferior frontal regions are connect by fibers of the corpus callosum (CC) (Hewitt, 1962). White matter fibers of the CC are among the most aligned fibers in the brain, connecting homotopic and heterotopic regions, and show high reliability in autoradiographic or MRI-based fiber tracking procedures (Kim *et al.*, 2008; Park *et al.*, 2008; Naets *et al.*, 2017). In terms of language development, the CC plays an important role for facilitating language lateralization (Hinkley *et al.*, 2016) as well the interplay between left and right inferior frontal areas for linguistic and paralinguistic aspects of speech processing as discussed above. Pathological changes in CC structure or development, such as congenital agenesis of the CC negatively affect speech and language abilities in later life (Adibpour *et al.*, 2018; Siffredi *et al.*, 2018). Given this importance of the CC as a structure for facilitating language development and speech, comparatively little in-vivo studies on interhemispheric language-related white matter pathways have been performed.

Much of the available tracking studies with diffusion tensor imaging (DTI) investigating interhemispheric connectivity, however, are not grounded in or related to neurolinguistics research but look at more general patterns of CC connections often based on deterministic tractography. A further methodological limitation of these most commonly used DTI methods **is that** the presence of crossing fibers often prevents a detailed mapping of lateral projections, such as the corpus callosum (Tuch *et al.*, 2002). The aim of our DTI-study here, using a probabilistic fiber tracking method that is particularly robust against crossing fibers, is to map the interhemispheric white matter fiber network that facilitates the interaction between left and right inferior frontal cortex in the healthy brain.

To this end, we use results from an fMRI language experiment, in which the paradigm involved linguistic computations of different complexity, for mapping the interhemispheric transcallosal fiber network between left and right inferior frontal regions with probabilistic DTI-based fiber tracking. Because the seed points for the probabilistic tractography derive from a well-controlled neurolinguistic fMRI experiment, we can relate the tracking results to specific aspects of interhemispheric callosal interaction based on linguistic complexity.

We show that highly aligned transcallosal fibers connect both left and right anterior-inferior IFC (BA45, ventral BA 44) and left and right posterior superior IFC (dorsal BA 44) to facilitate the rapid and dynamic interplay between linguistic and paralinguistic features in processing language.

## Methods

### Selection of seed coordinates for fiber tracking

The seed coordinates for the DTI-based tractography experiment performed for the study presented here were derived from an fMRI study on phonological transformation by Peschke et al. (2012); see their paper for the full description of the study (and the fMRI paradigm). Briefly, subjects in this experiment had to overtly REPEAT or TRANSFORM particular pseudo words or pseudo noun phrases (NPs) in the MR scanner. Peschke et al. modeled the pseudo words after names of countries (e.g., “Doga” [engl. “Doga”] in analogy to “Kuba” [engl. “Cuba”]) and the pseudo NPs were modeled after monosyllabic German NPs (e.g., “der Mall” [engl. e.g. “the goll”] in analogy to “der Ball” [engl. “the ball”]).

In the REPEAT condition, subjects had to just repeat the pseudo word or pseudo NP and in the TRANSFORM condition, the pseudo countries had to be transformed into the corresponding pseudo language (e.g. “Doga” -> “Doganisch” [engl. “Doga” -> “Dogan”]) and the pseudo NP into their corresponding diminutive form (e.g. “der Mall” -> “das Mällchen” [engl. e.g. “the goll” -> “the little goll”]). Linguistically, the transformation of the pseudo words entails mostly *prosodic* changes (PROSODIC), i.e. stress (“Dóga” -> “Dogánisch”), and transforming the pseudo NP requires more complex, *segmental* and *morphosyntactic* (SEGMENTAL), changes (e.g., in “der Ball” -> “das Bällchen” the segment “-all” is substituted with “-äll” and the pronoun changes from “der” to “das”).

The procedure for defining the seed points was the same as described in **(Kellmeyer *et al.*, 2013)**. For our tracking experiment, we used the suprathreshold coordinates from the random effects group-level fMRI analysis of the contrast TRANSFORM > REPEAT in the SEGMENTAL TRANSFORMATION. We did not use the random effects analysis of REPEAT alone for tracking because previous experiments have already demonstrated the structural connectivity patterns in the context of repetition of pseudo words via dorsal and ventral temporo-frontal pathways (Saur *et al.*, 2008, 2010).

In the contrast of interest, we identified the peak voxel coordinate in MNI space and then transformed it to the native space of each subjects’ DTI data and enlarged the seed coordinate to a seed sphere with a radius of 4 mm (containing 33 voxels), **see Table 1 for a list of the seeds and Figure 1 for the SPM visualization**. For better demarcation in defining the seed regions we chose a different threshold (p<0.001, uncorrected) for the t-maps in SPM8 on the original fMRI data from the fMRI study by (Peschke *et al.*, 2012). Therefore, peak coordinates, cluster size and t-values partly differ from Peschke et al. (2012), which used a threshold of p<0.05, FDR-corrected for the whole brain and a cluster level of >10 voxels. We should point out, that the sphere (containing 33 voxels) from which each tracking started encompassed the coordinate voxels from the published version of the study by **(Peschke *et al.*, 2012)** in each case. Thus, slight differences in the peak coordinates from our tracking experiment and the published version of Peschke et al. (2012) should not be a major concern as the sphere (containing 33 voxels) from which the tracking was started encompasses both the original coordinates and the coordinates used here.

**Table 1.** Seed regions for the DTI-based fiber tracking experiment; Abbreviations: IFG=inferior frontal gyrus, L=left,, R=right, MNI=Montreal Neurological Institute (atlas of brain coordinates); *at p<0.001, uncorrected

| Condition / task<br>in fMRI<br>experiment | Region | Hemisphere | Cluster<br>size<br>(voxels) | Peak MNI coordinates |  |  |  |
| --- | --- | --- | --- | --- | --- | --- | --- |
|  |  |  |  | x | y | z | t-value* |
| Segmental<br>manipulation /<br>transform ><br>repeat | IFG, pars opercularis | L | 1778 | -48 | 12 | 27 | 9.38 |
|  | IFG, pars triangularis | L | 1778 | -45 | 39 | 9 | 8.26 |
|  | IFG, pars opercularis | R | 613 | 45 | 12 | 24 | 5.12 |
|  | IFG, pars triangularis | R | 613 | 45 | 36 | 12 | 6.01 |

**Figure 1.**
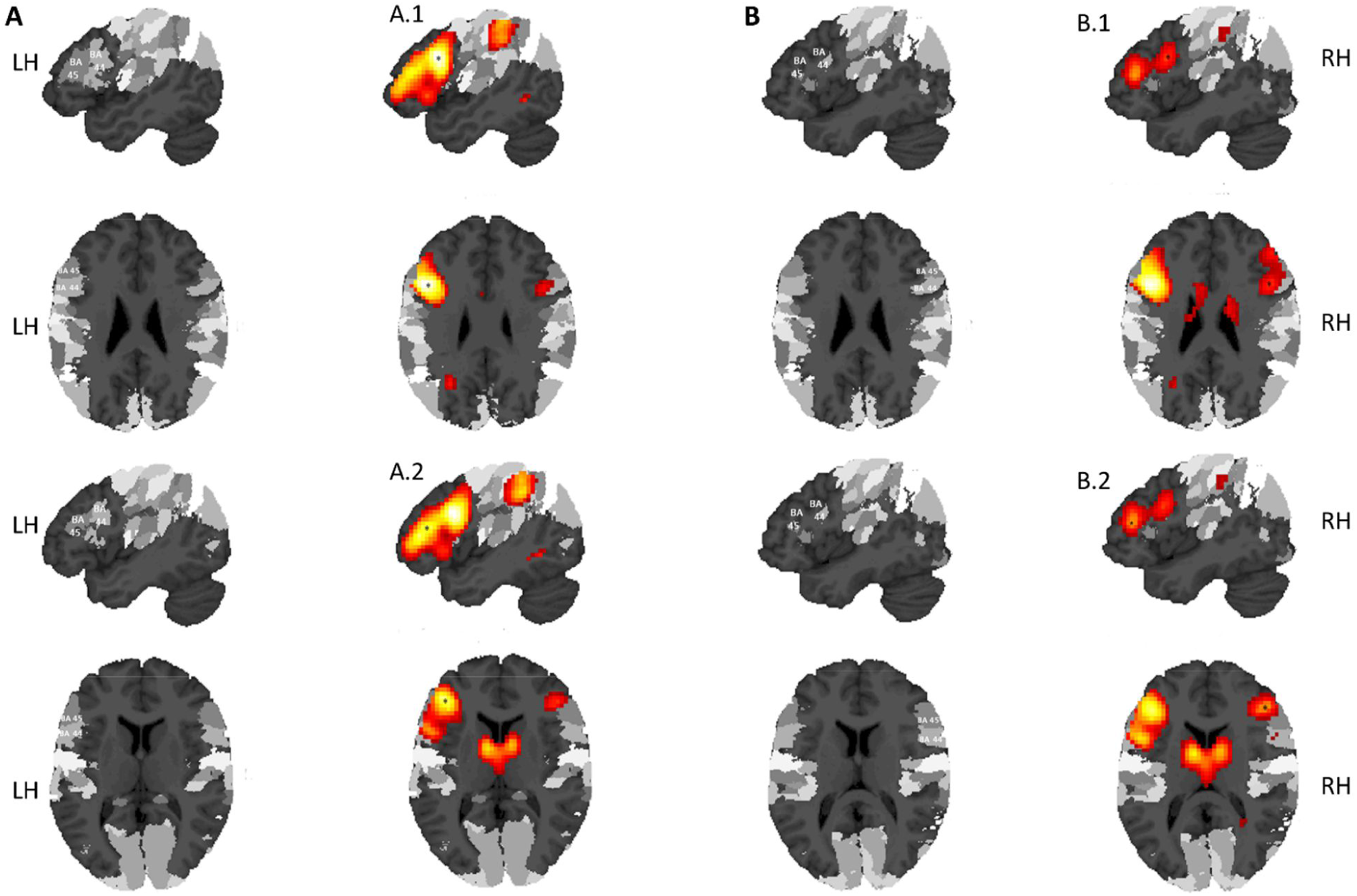
Suprathreshold peak coordinates (at p<0.001, uncorrected) in inferior frontal cortex that were used as seed coordinates for the interhemispheric probabilistic DTI-based fiber tracking are marked with an *. A1 shows the peak coordinate in LIFGpo, A2 in LIFGpt, B1 in RIFGpo, and B2 in RIFGpt. All clusters are superimposed on the cytoarchitectonic probability atlas by Eickhoff et al. (2005) in SPM8.

In the left hemisphere, we identified the two peak coordinates in the inferior frontal cortex (left inferior frontal gyrus [LIFG], pars opercularis [po], LIFG_po_; LIFG, pars triangularis [pt], LIFG_pt_) as seed coordinates. In the right hemisphere, we identified two frontal peaks (RIFG_po_; RIFG_pt_) as seed coordinates. **Table 1** and Fehler! Verweisquelle konnte nicht gefunden werden. provide an overview of the fMRI results showing the seed coordinates (A1, A2, B1, B2).

### Participants, group matching and methodological aspects of inter-group data analysis

The participants and the procedure for matching participants from the DTI group to the fMRI group (Peschke *et al.*, 2012) were the same as described in (Kellmeyer *et al.*, 2013).

In the study by (Peschke *et al.*, 2012), the researchers did not obtain DTI sequences from the participants in the fMRI study. For the DTI study presented here, we therefore matched twenty subjects in age, gender and handedness to the fMRI group. As in the fMRI study, all subjects were also native German speakers without any history of serious medical, neurological or psychiatric illness. All participants in this study were students recruited from the University of Freiburg and had therefore at least a diploma from German secondary school qualifying for university admission or matriculation as a basic level of education. We did not record information on socioeconomic status or employment record. Informed consent was obtained from all individual participants included in the study and all procedures performed in studies involving human participants were in accordance with the ethical standards of the institutional and/or national research committee and with the 1964 Helsinki declaration and its later amendments or comparable ethical standards. The DTI study was approved by the Ethics Committee of the Medical Center – University Medical **Center** of Freiburg.

The mean age in the DTI group was 24 years, the age range 20-38 years and 8 females and 12 males participated. Hand preference was tested with the 10-item version of the Edinburgh Handedness Inventory (Oldfield, 1971), subjects were identified as having predominantly right hand preference with an average laterality quotient of 0.8 (range 0.55-1.0). This group of participants did not differ significantly from the fMRI group in (Peschke *et al.*, 2012) in terms of age (p>0.1), gender (p>0.1) or handedness (p>0.1).

The matching of the groups should account for the majority of gross anatomical differences between two groups of healthy individuals, see also (Kellmeyer *et al.*, 2013; Suchan *et al.*, 2014). Furthermore, all subjects’ anatomical scans (T1) were checked for gross anatomical anomalies (if necessary with expertise from a qualified neuroradiologist).

#### Methodological aspects of inter-group and inter-method data collection and analysis

We use functional coordinates from one group of participants, the fMRI experiment by Peschke et al. (2012), for a fiber tracking experiment in another group of participants – in contrast to the widespread practice to perform functional and tracking experiments on the same group. We would however argue that our approach is not only legitimate but even preferable, considering the basic tenets of group-level statistical MRI-based data analysis. Generally, group-level neuroimaging data can be analyzed using either fixed-effects (FFX) or random-effects (RFX) analysis. While FFX analysis can be used for reporting (collections of multiple) case studies, the aim of RFX is to make statistical inferences about the population from which the group is drawn (Penny *et al.*, 2011). Today, RFX is widely used as standard approach in functional and structural neuroimaging data analysis, including the fMRI data set of the present study and had also been typically used in the functional data sets informing fiber tracking in other published reports. If, however, both (RFX) functional analysis and fiber tracking are performed in the same group of subjects, it is our view that it cannot be expected anymore that the resulting connectivity findings have any significance beyond the specific group that was investigated – i.e., inferences about the population from which the group was drawn become impossible, similar to FFX. Hence we used the functional localizers in a second group from the same population.

Given the group-level focus of our study here, we would argue that measuring and analyzing salient MRI data in two (carefully matched) groups of healthy participants should, if nothing else, increase the robustness and validity of our group level findings and inferences. These aspect are commensurate, in our view, with recent discussions in the neuroimaging community around establishing best practices and promoting open and reproducible measurement and analyses protocols in neuroimaging research (Nichols *et al.*, 2017; Poldrack *et al.*, 2017; Smith & Nichols, 2018).

We did not replicate the fMRI experiment from (Peschke et al., 2012) in the subjects of the DTI study here. While we agree that this would have been generally useful for validation of the fMRI results, we do not believe that it would have further improved the quality and validity of our tracking experiment. By transforming peak voxel coordinates into 33 voxel spheres as seed areas for the fiber tracking algorithm, we account for possible small inter-individual and inter-group differences in peak voxel localization, avoiding overspecification of individual peak voxel. Given limited resources on scanning time we therefore elected to solely perform the DTI measurements.

### DTI image acquisition

We acquired high angular resolution diffusion imaging (HARDI) data with a 3 Tesla *Siemens MAGNETOM Trio TIM* scanner using a diffusion-sensitive spin-echo echo planar imaging sequence with suppression of the cerebrospinal fluid signal. In total, we acquired 70 scans (with 69 slices) with 61 diffusion-encoding gradient directions (b-factor = 1000 s/mm) and 9 scans without diffusion weighting (b-factor = 0). The sequence parameters were: voxel size = 2×2×2 mm^3^, matrix size = 104×104 pixel^2^, TR = 11.8 s, TE = 96 ms, TI = 2.3 s. We corrected all scans for motion and distortion artifacts based on a reference measurement during reconstruction (Zaitsev *et al.*, 2004). Finally, we obtained a high-resolution T1 anatomical scan (160 slices, voxel size = 1×1×1 mm^3^, matrix=240×240 pixel^2^, TR=2.2 s, TE=2.6ms) for spatial processing of the DTI data.

### DTI-based probabilistic fiber tracking

We analyzed the DTI data by using the method of pathway extraction introduced by Kreher et al. (2008) which is part of the Matlab-based “DTI&Fiber toolbox” (Kreher *et al.*, 2008). This toolbox is available online for download (http://www.uniklinik-freiburg.de/mr/live/arbeitsgruppen/diffusion_en.html). Previously, this method has been used to identify white matter connections involved in language processing (Saur *et al.*, 2008, 2010; Kellmeyer *et al.*, 2013), attention (Umarova *et al.*, 2010) and motor cognition (Vry *et al.*, 2012; Hamzei *et al.*, 2015).

For this procedure, we first computed the effective self-diffusion tensor (sDT) from the HARDI dataset (Basser *et al.*, 1994), which was corrected for movement and distortion artifacts.

Then, we performed a Monte Carlo simulation of “random walks” to calculate the probabilistic maps for each seed region separately. This procedure is similar to the Probabilistic Index of Connectivity (PICo) method (Parker *et al.*, 2003). In extension to the PICo method, our probabilistic MCRW experiment preserves the information about the main traversing directions of the propagated fiber trajectories for each voxel. We then used this information for combining the probability maps. We extracted the orientation density function empirically from the effective diffusion tensor. The number of propagated trajectories was set to 10 and maximal fiber length was set to 150 voxels in accordance with our experience from the previous tracking studies mentioned above. We restricted the tracking area in each individual by a white matter mask to avoid tracking across anatomical borders. This mask included a small rim of grey matter to ensure that the cortical seed regions had indeed contact with the underlying white matter tracts.

To compute region-to-region anatomical connectivity between two seed spheres, we used a pairwise combination of two probability maps of interest (Kreher *et al.*, 2008). Computationally, this combination is a multiplication, which takes the main traversing trajectory of the random walk into account. Random walks starting from seed regions may face in either *opposing* directions (connecting fibers) or merge and face in the *same* direction (merging fibers). In a pathway connecting two seed regions, the proportion of connecting fibers should exceed the proportion of merging fibers. Using this directional information during the multiplication, merging fibers are suppressed and connecting fibers are preserved by the tracking algorithm (Kreher *et al.*, 2008). This procedure allows for extracting the most probable connecting pathway between two seed spheres without relying on *a priori* knowledge about the putative course of the white matter fibers. The resulting values represent a voxel-wise estimation of the probability that a particular voxel is part of the connecting fiber bundle of interest (represented by a “probability index forming part of the bundle of interest” [PIBI]). In order to extract the most probable fiber tracts connecting left and right inferior frontal regions, all left inferior frontal maps were combined permutatively with all right inferior frontal maps based on the respective linguistic context (prosodic or segmental manipulation).

### Post-processing of the individual probability maps

We scaled the individual probability maps to a range between 0 and 1. Then we spatially normalized the maps into standard Montreal Neurological Institute (MNI) space and subsequently smoothed them with an isotropic Gaussian kernel (3 mm) using SPM8. We computed *group maps* for each connection between seed regions by averaging the combined probability maps from all subjects. This resulted in one *mean group map* for each connection. Thus, any voxel in these group maps represents the arithmetic mean of the PIBI across subjects. To remove random artifacts, only voxels with PIBI values of >0.0145 were displayed, which excludes 95% of the voxels with PIBI >10^-6^. This cutoff value was empirically derived from the distribution observed in a large collection of preprocessed combined probability maps (Saur *et al.*, 2008). At the group level (n=20) we used a non-parametric statistic because PIBI values are not normally distributed (Saur *et al.*, 2010).

### Visualization and rendering of white matter fiber pathways

We visualized the resulting combined probability maps with the Matlab-based “DTI&Fiber Toolbox”, *MricroN* (http://www.sph.sc.edu/comd/rorden/mricron/) for 2D sections and rendered the fiber tracks with *OpenDX* by International Business Machines (IBM) (http://www.research.ibm.com/dx/).

## Results

The transcallosal fiber pathways between different subregions of left and right inferior frontal cortex show a homotopic region-to-region pattern of connectivity and the fiber systems are clearly segregated and aligned from a ventral anterior-inferior (left↔right IFG, partes triangularis) to dorsal posterior-superior (left↔right IFG, partes opercularis) gradient in the body and genu of the corpus callosum (Fig. 2). Importantly, the probabilistic tracking of the non-homotopic seed regions (i.e. A1-B2 and A2-B1 in Figure 2) did not yield significant group level anatomical connections. The observed pattern of clearly segregated and homotopic transcallosal pathways between L/R IFG_op_ and L/R IFG_pt_ was found in each individual participant of the studied group. The crossover tracking from left IFG_po_ to right IFG_pt_ and left IFG_pt_ to right IFG_po_, respectively, did not yield suprathreshold group level probability maps and are thus not visualized here.

**Figure 2.**
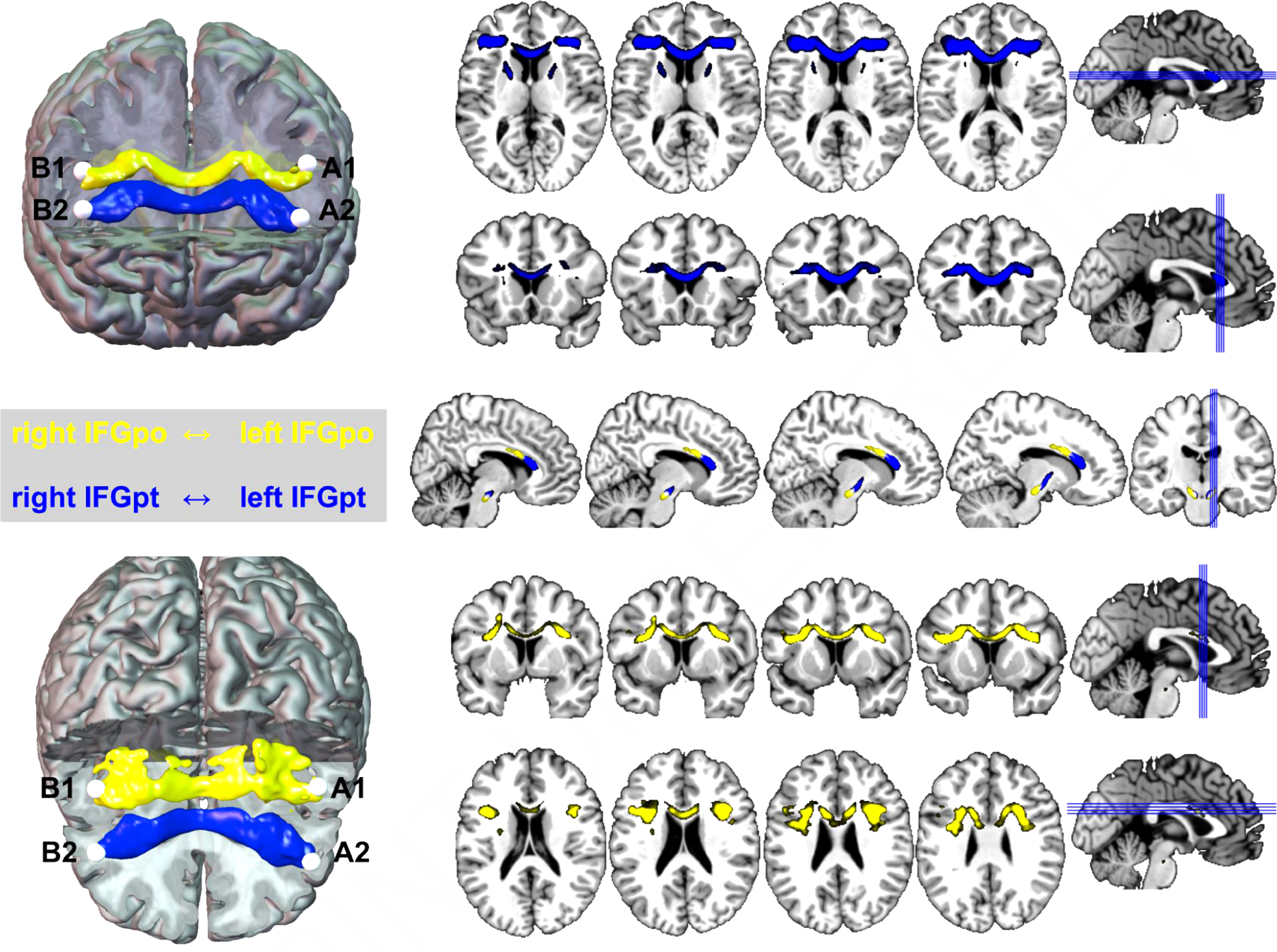
Mapping and rendering of the transcallosal white matter fiber tracts from left to right inferior frontal gyrus (pars opercularis, IFGpo, BA 44, in yellow) and left to right inferior frontal gyrus (pars triangularis, IFGpt, B45 and ventral BA44, in blue). Abbr.: A1, A2, B1, B2=seed coordinates from Fig.1.

## Discussion

The highly aligned transcallosal white matter pathways described here demonstrate a direct interhemispheric pathway for interaction between left and right homotopic inferior frontal cortex for language processing. The results are in agreement with previous anatomical *ex vivo* studies in humans on the topography of interhemispheric callosal fibers (Tomasch, 1954; Hewitt, 1962), as well as *in vivo* DTI-based parcellation studies (Huang *et al.*, 2005; Hofer & Frahm, 2006; Park *et al.*, 2008; Chao *et al.*, 2009; Teipel *et al.*, 2009; Fabri *et al.*, 2014). The results also show a close cross-species correspondence to homotopic patterns of interhemispheric fibers in non-human primates (Pandya *et al.*, 1971; Makris *et al.*, 2007; Phillips & Hopkins, 2012). Next, we first discuss the putative functional role of this transcallosal pathway connecting homotopic left and inferior frontal cortex for language processing in the healthy brain in the context of language development. Then, we analyze the potential role of this interhemispheric network for language reorganization and close with some remarks on comparative inter-species anatomy of the CC and its possible role for cognition.

### Function of interhemispheric inferior frontal connections for language development and function

In terms of functional significance, this transcallosal route may generally facilitate rapid neural processing for a variety of cognitive functions that involve homotopic inferior frontal regions in both hemispheres. In our own area of expertise - language function in the healthy and injured brain-it has long been recognized, that right and left inferior frontal cortex contribute substantially to paralinguistic aspects of language processing like prosody (George *et al.*, 1996; Hoekert *et al.*, 2010; Belyk & Brown, 2014) and/or speech rhythm (Riecker *et al.*, 2002; Geiser *et al.*, 2007; Jungblut *et al.*, 2012). More specifically, dysfunction of *intrinsic* (also called linguistic) features of prosody (such as stress, rhythm and pitch) seem to result from damage to *left* inferior frontal regions, whereas lesions to *right* inferior cortex often results in impaired processing of *extrinsic* features of speech like affective prosody – the tonal variation conveying emotions (Heilman *et al.*, 1984; Speedie *et al.*, 1984; Ross, 1993; Ross *et al.*, 1997; Belyk & Brown, 2014).

Relating this model of prosodic processing to the functional and structural interhemispheric inferior frontal network identified here, we propose the following interpretation: the transformation of the pseudo country into the respective pseudo language most prominently entails a shift of stress (“Dóga” -> “Dogánisch”)—a clear feature of linguistic prosody—but also the addition of segments and a change in grammatical category. This PROSODIC transformation condition (not shown in **Figure 1**, see Peschke et al. 2012) resulted in purely left fronto-parietal suprathreshold clusters, which is consistent with the putative role of LIFG in intrinsic / linguistic prosodic features of language in the models mentioned above. In the SEGMENTAL transformation task of the pseudo noun phrases (“Der Mall” -> “Das Mällchen”)—the basis of our DTI-tracking experiment here—more complex linguistic operations occur like morphosyntactic changes and a change of the pronoun. At the level of extrinsic / affective prosody, the features most often associated with the right IFC in the classic models, it would be difficult to construe different emotional valences of the pseudo-stimuli in the prosodic vs. the segmental transformation condition. Therefore, we interpret the involvement of right inferior frontal areas to reflect the greater *linguistic complexity* of the stimuli in the SEGMENTAL transformation condition. This is in accordance with previous fMRI studies and meta-analyses that have shown a right inferior frontal involvement in the context of complex linguistic processing independent of emotional content (Vigneau *et al.*, 2006, 2011; Price, 2010).

From a developmental perspective, the callosal transfer capabilities is also an important factor for the successful integration of linguistic and paralinguistic information in speech development. One DTI-based study, for example, showed that the macroanatomical thickness of the CC can be related to interhemispheric information transfer in a cohort of children between the age of six and eight (Westerhausen *et al.*, 2011) and another study in fifty-five children between the age of five and twelve showed that the diffusivity of callosal fibers—as a putative measure of fewer but larger callosal axons—is correlated with phonological skills (Dougherty *et al.*, 2007).

One important open question—which we cannot answer in the context of the study here—is, whether the functional role of interhemispheric callosal fibers in language processing is predominantly excitatory or inhibitory (Reggia *et al.*, 2001; Bloom & Hynd, 2005; van der Knaap & van der Ham, 2011). The clinical literature allows only for very limited and preliminary conclusions with regard to this question. Only very few clinical reports, let alone systematic studies, on the consequences for language processing of direct damage to callosal fibers connecting left and right inferior frontal cortex are available. Mostly, callosal dysfunction in relation to language was investigated in the context of congenital callosal disorders like agenesis or dysgenesis of the CC (Jeeves & Temple, 1987; Sanders, 1989; Paul *et al.*, 2003a; Brown *et al.*, 2005; Genç *et al.*, 2015). These studies have identified deficits in phonological and syntactic processing in relation to congenital CC dysfunction (Jeeves & Temple, 1987; Sanders, 1989; Temple *et al.*, 1989; Paul *et al.*, 2003b). Direct non-developmental callosal damage is a rare clinical event, mostly as a result of ischemic stroke or cerebral vasculitis (Mahale *et al.*, 2016). With respect to subsequent language dysfunction, we found only one report that describes a patient with a hemorrhagic lesion of the anterior part of the corpus callosum without damage to cortical projection areas (Klouda *et al.*, 1988). The patient initially showed a complete aprosody, that is a lack of tonal variation and reduced speed of speech, which recovered substantially throughout the follow-up period of one year.

From this body of clinical studies too, it is difficult to infer whether callosal fibers predominantly have an inhibitory or excitatory role in the transfer of linguistic computations in the brain. Next, we therefore highlight evidence from neuroimaging and non-invasive neurostimulation studies and research on language reorganization to shed light on the putative role of callosal fibers.

Other limitations for interpreting the results are: (1) Due to the relatively small sample size, analyzing the data for differences in sex-related interhemispheric white matter connectivity was not feasible in this study but would certainly be worthwhile in a larger sample of DTI data; (2) The interpretation of the findings here is also limited by the homogeneity of the subjects in terms of age (young) and education (university students), given that other demographic factors, such as socioeconomic status (especially during infancy / adolescence), may influence language development and skills (and the underlying neural network architecture); (3) Given the young average age of the subjects investigated here, the patterns of structural connectivity at a smaller scale might be different in the ageing brain, for example reflecting age-related reorganization of large-scale interhemispheric networks that support language processing.

Bearing these limitations in mind, the inferior frontal interhemispheric network described here may have important implications for language reorganization in the ageing brain and following injury, e.g. stroke.

### The role of inferior frontal callosal connections for language reorganization

FMRI studies on the dynamics of language reorganization have shown increased activity of right-hemispheric inferior frontal region following left hemispheric stroke in homologous inferior frontal cortex with post-stroke aphasia which has been interpreted as a sign of adaptive plasticity (Saur *et al.*, 2006; Winhuisen *et al.*, 2007). Importantly, the early up-regulation in the right hemisphere occurs in homotopic regions to the left hemispheric injured region which supports an important role of callosal fibers connecting homotopic regions (Staudt *et al.*, 2002). Further evidence corroborating the concept of right hemispheric involvement in aphasia recovery comes from research applying rTMS-based continuous theta burst stimulation (cTBS) over the left inferior frontal gyrus (Hartwigsen *et al.*, 2013). In this study, the virtual lesion of left inferior frontal cortex with cTBS resulted in up-regulation of the homotopic right inferior frontal cortex in reaction to the perturbation (Hartwigsen, 2015).The integrity of the interhemispheric white matter fiber network may therefore be critical to allow for adaptive recovery based on cross-hemispheric transfer. These studies on actual aphasia recovery and virtual lesion modelling support the concept of interhemispheric *inhibition* as the main role of homotopic inferior frontal callosal fibers (Bloom & Hynd, 2005; Kano *et al.*, 2012). This concept opens avenues for further exploring new concepts for stroke recovery with non-invasive and (invasive) neurostimulation as an emerging therapeutic approach (Hamilton *et al.*, 2011; Balossier *et al.*, 2015; Cherney, 2015; Otal *et al.*, 2015; Borich *et al.*, 2016).

To summarize, we interpret the homotopic interhemispheric inferior frontal white matter pathways, which we found here, as follows: the callosal connections of homotopic inferior frontal regions most likely supports integration of different levels of linguistic complexity. In this model, left inferior frontal regions would be sufficient to support linguistic transformations based on basic prosodic changes (like a shift of stress), whereas more complex linguistic operations, like segmental changes, additionally tap right inferior frontal regions for complementary computations. However, as real-time language comprehension and production requires fast information transfer for integration, it may be that this model of cooperative hierarchical processing of left and right IFC prefers homotopic regions. Processing of paralinguistic features like emotional prosody, which is less time-sensitive than computing linguistic features, in turn, may not depend upon strict homotopic connections. This model accounts for the hemispheric adaptive patterns in stroke recovery discussed above, as well as the neurotypology of disorders of prosodic perception and production as a result of right hemispheric inferior frontal injury (Blonder *et al.*, 1991; Hoekert *et al.*, 2010; Belyk & Brown, 2014). Finally, as this study is an in-vivo anatomical study based on DTI, we want to highlight briefly some interesting inter-species features of the anatomy and function of the CC.

### Some remarks on interspecies anatomy and function of the corpus callosum

From an evolutionary and mammalian inter-species perspective, the CC is a highly conserved macroanatomic structure, indicating that it is important for supporting a variety of interhemispheric computations, independent of (but in humans including) language processing (Olivares *et al.*, 2000, 2001; Aboitiz & Montiel, 2003). A MRI tractography study in chimpanzees, the closest species related to humans which has been studied with DTI to date, shows a very similar topical pattern of interhemispheric connections and fiber alignment in the CC as human tractography studies and our results here(Phillips & Hopkins, 2012). If we move further away back on the eutherian clade of our evolutionary ancestry, however, fine differences in CC microstructure and connectivity emerge. In the largest cross-species study to date, Olivares et al. (Olivares *et al.*, 2001) demonstrated that the proportional numeric composition of fibers of the CC is preserved across six different species (the rat, the rabbit, the cat, the dog, the horse and the cow). Whereas the number of callosal fibers does not scale with increased brain size, the fiber diameter (and hence conduction velocity) does. This indicates that the type of fiber and quite likely also the pattern of connectivity might determine interhemispheric information transfer capabilities and that callosal transmission time may not be constant across species. These fine differences in interhemispheric network architecture and conduction properties, in turn, may relate to differences in the cognitive abilities of different mammalian species through processing constraints (Aboitiz *et al.*, 2003). For humans this structurally and functionally honed interplay and division of labor between the left and right hemisphere may indeed be the prerequisite for complex cognition and, ultimately, “the human condition” (Gazzaniga, 2000).

## Acknowledgments

The authors would like to thank Professor Dorothee Saur (University of Leipzig – Medical Center) for valuable discussions of the work presented here. *Funding*: This work was (partly) supported by the German Ministry of Education and Research (BMBF) (grant number 13GW0053D) to the University of Freiburg – Medical Center, and the German Research Foundation (DFG) (grant number EXC1086) to the University of Freiburg, Germany. *Other:* Parts of this study were used for the dissertation of author P.K. (University of Freiburg, Faculty of Medicine).

## Statement regarding potential conflicts of interest

The authors have no conflict of interest to declare.

## Author contributions

Philipp Kellmeyer: data collection, experimental design, data analysis, writing of the manuscript Magnus-Sebastian Vry: experimental design, data analysis, writing of the manuscript Tonio Ball: experimental design, reviewing of the manuscript.

## Data accessibility

The datasets analyzed for this study are available from the corresponding on reasonable request.

## Abbreviations

BA: Brodmann Area
CC: corpus callosum
DTI: diffusion tensor imaging
HARDI: high angular resolution diffusion imaging [a DTI measurement technique]
IBM: International Business Machines
IFC: inferior frontal cortex
IFG: inferior frontal gyrus
IFGpo: inferior frontal gyrus pars opercularis
IFGpt: inferior frontal gyrus pars triangularis
FFX: fixed effect [in analyzing fMRI data]
fMRI: functional magnetic resonance imaging
L: left
MCSRW: Monte Carlo simulation of random walks [a statistical method]
mm: millimeter
MNI: Montreal Neurological Institute
MRI: magnetic resonance imaging
ms: milliseconds
PIBI: probability index forming part of the bundle of interest
PICo: Probabilistic Index of Connectivity
R: right
RFX: random effects [in analyzing fMRI data]
rTMS: repetitive transcranial magnetic stimulation
s: seconds
SPM: statistical parametric mapping
TE: echo time [a specific MRI pulse sequence parameter]
TR: repetition time [a specific MRI pulse sequence parameter]

## References

Aboitiz, F., López, J., & Montiel, J. (2003) Long distance communication in the human brain: timing constraints for inter-hemispheric synchrony and the origin of brain lateralization. Biol. Res., 36, 89–99.

Aboitiz, F. & Montiel, J. (2003) One hundred million years of interhemispheric communication: the history of the corpus callosum. Braz. J. Med. Biol. Res., 36, 409–420.

Adibpour, P., Dubois, J., Moutard, M.-L., & Dehaene-Lambertz, G. (2018) Early asymmetric inter-hemispheric transfer in the auditory network: insights from infants with corpus callosum agenesis. Brain Struct. Funct., 223, 2893–2905.

Amunts, K., Schleicher, A., Bürgel, U., Mohlberg, H., Uylings, H.B.M., & Zilles, K. (1999) Broca’s region revisited: Cytoarchitecture and intersubject variability. J. Comp. Neurol., 412, 319–341.

Balossier, A., Etard, O., Descat, C., Vivien, D., & Emery, E. (2015) Epidural Cortical Stimulation as a Treatment for Poststroke Aphasia A Systematic Review of the Literature and Underlying Neurophysiological Mechanisms. Neurorehabil. Neural Repair, 1545968315606989.

Basser, P.J., Mattiello, J., & LeBihan, D. (1994) Estimation of the effective self-diffusion tensor from the NMR spin echo. J. Magn. Reson. B, 103, 247–254.

Belyk, M. & Brown, S. (2014) Perception of affective and linguistic prosody: an ALE meta-analysis of neuroimaging studies. Soc. Cogn. Affect. Neurosci., 9, 1395–1403.

Blonder, L.X., Bowers, D., & Heilman, K.M. (1991) The Role of the Right Hemisphere in Emotional Communication. Brain, 114, 1115–1127.

Bloom, J.S. & Hynd, G.W. (2005) The Role of the Corpus Callosum in Interhemispheric Transfer of Information: Excitation or Inhibition? Neuropsychol. Rev., 15, 59–71.

Bookheimer, S. (2002) FUNCTIONAL MRI OF LANGUAGE: New Approaches to Understanding the Cortical Organization of Semantic Processing. Annu. Rev. Neurosci., 25, 151–188.

Borich, M.R., Wheaton, L.A., Brodie, S.M., Lakhani, B., & Boyd, L.A. (2016) Evaluating interhemispheric cortical responses to transcranial magnetic stimulation in chronic stroke: A TMS-EEG investigation. Neurosci. Lett., 618, 25–30.

Bornkessel-Schlesewsky, I. & Schlesewsky, M. (2013) Reconciling time, space and function: A new dorsal–ventral stream model of sentence comprehension. Brain Lang., 125, 60–76.

Brown, W.S., Symingtion, M., VanLancker-Sidtis, D., Dietrich, R., & Paul, L.K. (2005) Paralinguistic processing in children with callosal agenesis: Emergence of neurolinguistic deficits. Brain Lang., 93, 135–139.

Chao, Y.-P., Cho, K.-H., Yeh, C.-H., Chou, K.-H., Chen, J.-H., & Lin, C.-P. (2009) Probabilistic topography of human corpus callosum using cytoarchitectural parcellation and high angular resolution diffusion imaging tractography. Hum. Brain Mapp., 30, 3172–3187.

Cherney, L.R. (2015) Epidural Cortical Stimulation as Adjunctive Treatment for Nonfluent Aphasia Phase 1 Clinical Trial Follow-up Findings. Neurorehabil. Neural Repair, 1545968315622574.

Dapretto, M. & Bookheimer, S.Y. (1999) Form and Content: Dissociating Syntax and Semantics in Sentence Comprehension. Neuron, 24, 427–432.

Dougherty, R.F., Ben-Shachar, M., Deutsch, G.K., Hernandez, A., Fox, G.R., & Wandell, B.A. (2007) Temporal-callosal pathway diffusivity predicts phonological skills in children. Proc. Natl. Acad. Sci., 104, 8556–8561.

Fabri, M., Pierpaoli, C., Barbaresi, P., & Polonara, G. (2014) Functional topography of the corpus callosum investigated by DTI and fMRI. World J. Radiol., 6, 895–906.

Frühholz, S., Gschwind, M., & Grandjean, D. (2015) Bilateral dorsal and ventral fiber pathways for the processing of affective prosody identified by probabilistic fiber tracking. NeuroImage, 109, 27–34.

Gazzaniga, M.S. (2000) Cerebral specialization and interhemispheric communication. Brain, 123, 1293–1326.

Geiser, E., Zaehle, T., Jancke, L., & Meyer, M. (2007) The Neural Correlate of Speech Rhythm as Evidenced by Metrical Speech Processing. J. Cogn. Neurosci., 20, 541–552.

Genç, E., Ocklenburg, S., Singer, W., & Güntürkün, O. (2015) Abnormal interhemispheric motor interactions in patients with callosal agenesis. Behav. Brain Res., 293, 1–9.

George, M.S., Parekh, P.I., Rosinsky, N., Ketter, T.A., Kimbrell, T.A., Heilman, K.M., Herscovitch, P., & Post, R.M. (1996) Understanding Emotional Prosody Activates Right Hemisphere Regions. Arch. Neurol., 53, 665–670.

Hamilton, R.H., Chrysikou, E.G., & Coslett, B. (2011) Mechanisms of Aphasia Recovery After Stroke and the Role of Noninvasive Brain Stimulation. Brain Lang., 118, 40–50.

Hamzei, F., Vry, M.-S., Saur, D., Glauche, V., Hoeren, M., Mader, I., Weiller, C., & Rijntjes, M. (2015) The Dual-Loop Model and the Human Mirror Neuron System: an Exploratory Combined fMRI and DTI Study of the Inferior Frontal Gyrus. Cereb. Cortex, bhv066.

Hartwigsen, G. (2015) The neurophysiology of language: Insights from non-invasive brain stimulation in the healthy human brain. Brain Lang., The electrophysiology of speech, language, and its precursors, 148, 81–94.

Hartwigsen, G., Saur, D., Price, C.J., Ulmer, S., Baumgaertner, A., & Siebner, H.R. (2013) Perturbation of the left inferior frontal gyrus triggers adaptive plasticity in the right homologous area during speech production. Proc. Natl. Acad. Sci., 110, 16402–16407.

Heilman, K.M., Bowers, D., Speedie, L., & Coslett, H.B. (1984) Comprehension of affective and nonaffective prosody. Neurology, 34, 917–921.

Hewitt, W. (1962) The development of the human corpus callosum. J. Anat., 96, 355–358.

Hillis, A.E. (2007) Aphasia Progress in the last quarter of a century. Neurology, 69, 200–213.

Hinkley, L.B.N., Marco, E.J., Brown, E.G., Bukshpun, P., Gold, J., Hill, S., Findlay, A.M., Jeremy, R.J., Wakahiro, M.L., Barkovich, A.J., Mukherjee, P., Sherr, E.H., & Nagarajan, S.S. (2016) The Contribution of the Corpus Callosum to Language Lateralization. J. Neurosci., 36, 4522–4533.

Hoekert, M., Vingerhoets, G., & Aleman, A. (2010) Results of a pilot study on the involvement of bilateral inferior frontal gyri in emotional prosody perception: an rTMS study. BMC Neurosci., 11, 93.

Hofer, S. & Frahm, J. (2006) Topography of the human corpus callosum revisited—Comprehensive fiber tractography using diffusion tensor magnetic resonance imaging. NeuroImage, 32, 989–994.

Huang, H., Zhang, J., Jiang, H., Wakana, S., Poetscher, L., Miller, M.I., van Zijl, P.C., Hillis, A.E., Wytik, R., & Mori, S. (2005) DTI tractography based parcellation of white matter: Application to the mid-sagittal morphology of corpus callosum. NeuroImage, 26, 195–205.

Jeeves, M.A. & Temple, C.M. (1987) A further study of language function in callosal agenesis. Brain Lang., 32, 325–335.

Jungblut, M., Huber, W., Pustelniak, M., & Schnitker, R. (2012) The impact of rhythm complexity on brain activation during simple singing: An event-related fMRI study. Restor. Neurol. Neurosci., 30.

Kano, T., Kobayashi, M., Ohira, T., & Yoshida, K. (2012) Speech-induced modulation of interhemispheric inhibition. Neurosci. Lett., 531, 86–90.

Kellmeyer, P., Ziegler, W., Peschke, C., Juliane, E., Schnell, S., Baumgaertner, A., Weiller, C., & Saur, D. (2013) Fronto-parietal dorsal and ventral pathways in the context of different linguistic manipulations. Brain Lang., 127, 241–250.

Kim, E.Y., Park, H.-J., Kim, D.-H., Lee, S.-K., & Kim, J. (2008) Measuring Fractional Anisotropy of the Corpus Callosum Using Diffusion Tensor Imaging: Mid-Sagittal versus Axial Imaging Planes. Korean J. Radiol., 9, 391–395.

Klouda, G.V., Robin, D.A., Graff-Radford, N.R., & Cooper, W.E. (1988) The role of callosal connections in speech prosody. Brain Lang., 35, 154–171.

Kreher, B.W., Schnell, S., Mader, I., Il’yasov, K.A., Hennig, J., Kiselev, V.G., & Saur, D. (2008) Connecting and merging fibres: Pathway extraction by combining probability maps. NeuroImage, 43, 81–89.

Levy, J. & Trevarthen, C. (1977) Perceptual, semantic and phonetic aspects of elementary language processes in split-brain patients. Brain J. Neurol., 100 Pt 1, 105–118.

Mahale, R., Mehta, A., Buddaraju, K., John, A.A., Javali, M., & Srinivasa, R. (2016) Diffuse corpus callosum infarction — Rare vascular entity with differing etiology. J. Neurol. Sci., 360, 45–48.

Makris, N., Papadimitriou, G.M., van der Kouwe, A., Kennedy, D.N., Hodge, S.M., Dale, A.M., Benner, T., Wald, L.L., Wu, O., Tuch, D.S., Caviness, V.S., Moore, T.L., Killiany, R.J., Moss, M.B., & Rosene, D.L. (2007) Frontal connections and cognitive changes in normal aging rhesus monkeys: a DTI study. Neurobiol. Aging, 28, 1556–1567.

Monrad-Krohn, G.H. (1947) The Prosodic Quality of Speech and Its Disorders: (a Brief Survey from a Neurologist’s Point of View). Acta Psychiatr. Scand., 22, 255–269.

Monrad-Krohn, G.H. (1957) The Third Element of Speech: Prosody in the Neuro-Psychiatric Clinic. Br. J. Psychiatry, 103, 326–331.

Naets, W., Loon, J.V., Paglioli, E., Paesschen, W.V., Palmini, A., & Theys, T. (2017) Callosotopy: leg motor connections illustrated by fiber dissection. Brain Struct. Funct., 222, 661–667.

Nichols, T.E., Das, S., Eickhoff, S.B., Evans, A.C., Glatard, T., Hanke, M., Kriegeskorte, N., Milham, M.P., Poldrack, R.A., Poline, J.-B., Proal, E., Thirion, B., Van Essen, D.C., White, T., & Yeo, B.T.T. (2017) Best practices in data analysis and sharing in neuroimaging using MRI. Nat. Neurosci., 20, 299–303.

Oldfield, R.C. (1971) The assessment and analysis of handedness: the Edinburgh inventory. Neuropsychologia, 9, 97–113.

Olivares, R., Michalland, S., & Aboitiz, F. (2000) Cross-Species and Intraspecies Morphometric Analysis of the Corpus Callosum. Brain. Behav. Evol., 55, 37–43.

Olivares, R., Montiel, J., & Aboitiz, F. (2001) Species Differences and Similarities in the Fine Structure of the Mammalian Corpus callosum. Brain. Behav. Evol., 57, 98–105.

Otal, B., Olma, M.C., Flöel, A., & Wellwood, I. (2015) Inhibitory non-invasive brain stimulation to homologous language regions as an adjunct to speech and language therapy in post-stroke aphasia: a meta-analysis. Front. Hum. Neurosci., 9.

Pandya, D.N., Karol, E.A., & Heilbronn, D. (1971) The topographical distribution of interhemispheric projections in the corpus callosum of the rhesus monkey. Brain Res., 32, 31–43.

Park, H.-J., Kim, J.J., Lee, S.-K., Seok, J.H., Chun, J., Kim, D.I., & Lee, J.D. (2008) Corpus callosal connection mapping using cortical gray matter parcellation and DT-MRI. Hum. Brain Mapp., 29, 503–516.

Parker, G.J.M., Haroon, H.A., & Wheeler-Kingshott, C.A.M. (2003) A framework for a streamline-based probabilistic index of connectivity (PICo) using a structural interpretation of MRI diffusion measurements. J. Magn. Reson. Imaging JMRI, 18, 242–254.

Paul, L.K., Van Lancker-Sidtis, D., Schieffer, B., Dietrich, R., & Brown, W.S. (2003a) Communicative deficits in agenesis of the corpus callosum: Nonliteral language and affective prosody. Brain Lang., 85, 313–324.

Paul, L.K., Van Lancker-Sidtis, D., Schieffer, B., Dietrich, R., & Brown, W.S. (2003b) Communicative deficits in agenesis of the corpus callosum: Nonliteral language and affective prosody. Brain Lang., 85, 313–324.

Penny, W.D., Friston, K.J., Ashburner, J.T., Kiebel, S.J., & Nichols, T.E. (2011) Statistical Parametric Mapping: The Analysis of Functional Brain Images. Elsevier.

Peschke, C., Ziegler, W., Eisenberger, J., & Baumgaertner, A. (2012) Phonological manipulation between speech perception and production activates a parieto-frontal circuit. NeuroImage, Neuroergonomics: The human brain in action and at work, 59, 788–799.

Phillips, K.A. & Hopkins, W.D. (2012) Topography of the Chimpanzee Corpus Callosum. PLoS ONE, 7.

Poeppel, D., Emmorey, K., Hickok, G., & Pylkkänen, L. (2012) Towards a new neurobiology of language. J. Neurosci. Off. J. Soc. Neurosci., 32, 14125–14131.

Poldrack, R.A., Baker, C.I., Durnez, J., Gorgolewski, K.J., Matthews, P.M., Munafò, M.R., Nichols, T.E., Poline, J.-B., Vul, E., & Yarkoni, T. (2017) Scanning the horizon: towards transparent and reproducible neuroimaging research. Nat. Rev. Neurosci., advance online publication.

Price, C.J. (2010) The anatomy of language: a review of 100 fMRI studies published in 2009. Ann. N. Y. Acad. Sci., 1191, 62–88.

Price, C.J. (2012) A review and synthesis of the first 20 years of PET and fMRI studies of heard speech, spoken language and reading. NeuroImage, 20 YEARS OF fMRI20 YEARS OF fMRI, 62, 816–847.

Reggia, J.A., Goodall, S.M., Shkuro, Y., & Glezer, M. (2001) The callosal dilemma: Explaining diaschisis in the context of hemispheric rivalry via a neural network model. Neurol. Res., 23, 465–471.

Riecker, A., Wildgruber, D., Dogil, G., Grodd, W., & Ackermann, H. (2002) Hemispheric Lateralization Effects of Rhythm Implementation during Syllable Repetitions: An fMRI Study. NeuroImage, 16, 169–176.

Ross, E.D. (1993) Nonverbal aspects of language. Neurol. Clin., 11, 9–23.

Ross, E.D., Thompson, R.D., & Yenkosky, J. (1997) Lateralization of Affective Prosody in Brain and the Callosal Integration of Hemispheric Language Functions. Brain Lang., 56, 27–54.

Sammler, D., Grosbras, M.-H., Anwander, A., Bestelmeyer, P.E.G., & Belin, P. (2015) Dorsal and Ventral Pathways for Prosody. Curr. Biol., 25, 3079–3085.

Sanders, R.J. (1989) Sentence comprehension following agenesis of the corpus callosum. Brain Lang., 37, 59–72.

Saur, D., Kreher, B.W., Schnell, S., Kümmerer, D., Kellmeyer, P., Vry, M.-S., Umarova, R., Musso, M., Glauche, V., Abel, S., Huber, W., Rijntjes, M., Hennig, J., & Weiller, C. (2008) Ventral and dorsal pathways for language. Proc. Natl. Acad. Sci. U. S. A., 105, 18035–18040.

Saur, D., Lange, R., Baumgaertner, A., Schraknepper, V., Willmes, K., Rijntjes, M., & Weiller, C. (2006) Dynamics of language reorganization after stroke. Brain, 129, 1371–1384.

Saur, D., Schelter, B., Schnell, S., Kratochvil, D., Küpper, H., Kellmeyer, P., Kümmerer, D., Klöppel, S., Glauche, V., Lange, R., Mader, W., Feess, D., Timmer, J., & Weiller, C. (2010) Combining functional and anatomical connectivity reveals brain networks for auditory language comprehension. NeuroImage, 49, 3187–3197.

Shalom, D.B. & Poeppel, D. (2008) Functional Anatomic Models of Language: Assembling the Pieces. The Neuroscientist, 14, 119–127.

Siffredi, V., Anderson, V., McIlroy, A., Wood, A.G., Leventer, R.J., & Spencer-Smith, M.M. (2018) A Neuropsychological Profile for Agenesis of the Corpus Callosum? Cognitive, Academic, Executive, Social, and Behavioral Functioning in School-Age Children. J. Int. Neuropsychol. Soc., 24, 445–455.

Smith, S.M. & Nichols, T.E. (2018) Statistical Challenges in “Big Data” Human Neuroimaging. Neuron, 97, 263–268.

Speedie, L.J., Coslett, H.B., & Heilman, K.M. (1984) Repetition of Affective Prosody in Mixed Transcortical Aphasia. Arch. Neurol., 41, 268–270.

Staudt, M., Lidzba, K., Grodd, W., Wildgruber, D., Erb, M., & Krägeloh-Mann, I. (2002) Right-Hemispheric Organization of Language Following Early Left-Sided Brain Lesions: Functional MRI Topography. NeuroImage, 16, 954–967.

Suchan, J., Umarova, R., Schnell, S., Himmelbach, M., Weiller, C., Karnath, H.-O., & Saur, D. (2014) Fiber pathways connecting cortical areas relevant for spatial orienting and exploration. Hum. Brain Mapp., 35, 1031–1043.

Teipel, S.J., Pogarell, O., Meindl, T., Dietrich, O., Sydykova, D., Hunklinger, U., Georgii, B., Mulert, C., Reiser, M.F., Möller, H.-J., & Hampel, H. (2009) Regional networks underlying interhemispheric connectivity: An EEG and DTI study in healthy ageing and amnestic mild cognitive impairment. Hum. Brain Mapp., 30, 2098–2119.

Temple, C.M., Jeeves, M.A., & Vilarroya, O. (1989) Ten pen men: rhyming skills in two children with callosal agenesis. Brain Lang., 37, 548–564.

Tomasch, J. (1954) Size, distribution, and number of fibres in the human corpus callosum. Anat. Rec., 119, 119–135.

Tuch, D.S., Reese, T.G., Wiegell, M.R., Makris, N., Belliveau, J.W., & Wedeen, V.J. (2002) High angular resolution diffusion imaging reveals intravoxel white matter fiber heterogeneity. Magn. Reson. Med., 48, 577–582.

Umarova, R.M., Saur, D., Schnell, S., Kaller, C.P., Vry, M.-S., Glauche, V., Rijntjes, M., Hennig, J., Kiselev, V., & Weiller, C. (2010) Structural Connectivity for Visuospatial Attention: Significance of Ventral Pathways. Cereb. Cortex, 20, 121–129.

van der Knaap, L.J. & van der Ham, I.J.M. (2011) How does the corpus callosum mediate interhemispheric transfer? A review. Behav. Brain Res., 223, 211–221.

Vigneau, M., Beaucousin, V., Hervé, P.Y., Duffau, H., Crivello, F., Houdé, O., Mazoyer, B., & Tzourio-Mazoyer, N. (2006) Meta-analyzing left hemisphere language areas: Phonology, semantics, and sentence processing. NeuroImage, 30, 1414–1432.

Vigneau, M., Beaucousin, V., Hervé, P.-Y., Jobard, G., Petit, L., Crivello, F., Mellet, E., Zago, L., Mazoyer, B., & Tzourio-Mazoyer, N. (2011) What is right-hemisphere contribution to phonological, lexico-semantic, and sentence processing?: Insights from a meta-analysis. NeuroImage, 54, 577–593.

Vry, M.-S., Saur, D., Rijntjes, M., Umarova, R., Kellmeyer, P., Schnell, S., Glauche, V., Hamzei, F., & Weiller, C. (2012) Ventral and dorsal fiber systems for imagined and executed movement. Exp. Brain Res., 219, 203–216.

Westerhausen, R., Luders, E., Specht, K., Ofte, S.H., Toga, A.W., Thompson, P.M., Helland, T., & Hugdahl, K. (2011) Structural and Functional Reorganization of the Corpus Callosum between the Age of 6 and 8 Years. Cereb. Cortex, 21, 1012–1017.

Winhuisen, L., Thiel, A., Schumacher, B., Kessler, J., Rudolf, J., Haupt, W.F., & Heiss, W.D. (2007) The Right Inferior Frontal Gyrus and Poststroke Aphasia A Follow-Up Investigation. Stroke, 38, 1286–1292.

Zaitsev, M., Hennig, J., & Speck, O. (2004) Point spread function mapping with parallel imaging techniques and high acceleration factors: fast, robust, and flexible method for echo-planar imaging distortion correction. Magn. Reson. Med., 52, 1156–1166.

